# On the Geometry of Interacting Axon Tracts in Relation to Their Functional Properties

**DOI:** 10.1101/2022.03.29.486317

**Authors:** Aman Chawla, Salvatore Domenic Morgera

## Abstract

It is well known that demyelination associated with multiple sclerosis alters coupling between axons in a diseased nerve tract due to exposure of ion channels. In this work we ask the question whether the presence of electric fields as a coupling mechanism between nerve axons, has any impact on the expected conduction properties of the nerve under focal demyelination. Using an approximate geometry for a bundle of three axons, we allow electric fields generated at a node of Ranvier to influence other nodes of Ranvier on different axons. In order to introduce focal demyelination, we allow a new type of structure, intermediate between a node and an internodal segment. We find that the nerve’s conduction profiles are different with and without field-based coupling. This is important because it opens up the possibility for an experimental investigation to study whether field-based coupling predominates over current-based coupling. Fields and currents may also be measured using different methods and so there are implications for bio-instrumentation. From a systems perspective, the study is based on a novel equation which allows electric fields to act as sources of current at axons’ nodes of Ranvier, thereby generating new behaviors from the set of coupled partial differential equations that model the electrodynamics of the nerve. In addition to our simulation study of focal demyelination, we present a justification for a certain geometric *W*-matrix used in our phenomenological 2019 model. We also provide the first simulations of a tract of tortuous axons, showing the impact of tract tortuosity on conduction velocity. The tortuous model is analyzed geometrically to illustrate the mechanism of synchronization. Finally, this same model is shown to be castable as a learning machine - a formulation that has neuroscience implications.

## 1 Introduction

Focal demyelination has been studied extensively in the biological literature. In [1], the authors look at the corpus callosum of the rabbit and find a way to induce focal demyelination. In [2], death of oligodendrocytes was associated with focal demyelination in the adult rat brain, and the study in [3] uses novel MRI techniques to image focal demyelination in vivo induced by lysolecithin in male Lewis rats.

Numerical methods in the neurosciences have a long history dating at least to Hodgkin and Huxley [4]. While we celebrate the remarkable success of Hodgkin-Huxley neural theory [5] in mimicking the real nerve, there is still no full analytical solution to those equations; they were initially partly composed on the basis of empirical fits to observed data. This is a constraint on our understanding of nerve function. One would ideally need to delve into ion channel structure and function and perhaps even incorporate the molecular-quantum level and then step upward classically to get an actual ‘derivation’ of the Hodgkin Huxley formula from first principles. This is undoubtedly a hard task and has not been accomplished by anyone yet.

From the mathematical and biophysical perspective, studies on focal demyelination are thus relatively few. Papers such as [6], within the past half-century, have looked at demyelination in the context of stand-alone axons, providing computer simulation results. In the present paper, we focus on current and field-coupled axon tracts as the arena in which to carry out this investigation. Another paper, [7], also looks at focal demyelination. However, in it the authors vary the paranodal seal resistance, whereas our work is carried out on the basis of varying the paranodal capacitance. They have regularly spaced paranodes throughout their model, whereas we topically insert paranodes to test their impact. Apart from these differences, they do not consider coupling between axons, either via currents or fields, which we consider in this paper. The other papers of the same group, such as [8], follow their 1995 model and hence the differences from our work carry over to these later papers as well.

Note that current-mediated coupling (as in the term ‘current-coupled’) is commonly known as ephaptic coupling in the literature and is based on a return path of the axoplasmic current through the extracellular space [9]. Field-mediated coupling (as in the term ‘field-coupled’) as discussed in the present paper allows the endogenous electric fields generated at nodes of Ranvier to influence the movement of and injection of ions into distant nodes of Ranvier on different axons and doesn’t have an exact parallel in the literature. However, for a comparison, see [10], where the phrase ‘ephaptic interactions’ is used in a somewhat similar context, with the mention of ‘fields’. Their Equations (4) and (5) however, appear to be a recasting of Equations (9) and (11) of [9].

In the present paper we will take the Hodgkin-Huxley formulation for granted, and build on top of it an understanding of demyelination, showing an important application of the program to develop a deeper mathematical and biophysical understanding of axon-axon coupling. Towards this end, the paper is structured as follows. First, in Section 2 we describe the model used in this study and present a working example. We also elucidate an important part of the model which was phenomenologically justified previously. We provide a geometric analysis of synchronization and frame the tortuous tract as a learning machine. Thereafter, in Section 3 we study focal demyelination in axon tracts under current-mediated coupling as well as under field-mediated coupling, also providing simulations of conduction under tortuosity. Finally, we conclude in Section 4. The appendix looks at the geometry of the tract, as encoded in the tract matrix, in the sense of information theory and illuminates a perspective on the progression of multiple sclerosis that has not been considered before.

## 2 Materials and Methods

### 2.1 Mathematical Model and Its Justification

The mathematical model that we use in this paper is described in [11]. The central equation is presented here as well for a self-contained treatment. The notation is succinctly described in Table 1.

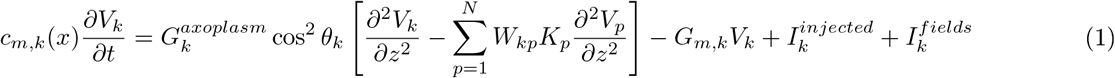

**Table 1:** Notational Summary

| Symbol | Meaning |
| --- | --- |
| $c_{m,k}$ | per unit length capacitance of the node, paranode or internode |
| $V, V_k$ | transmembrane voltage of the ($k$-th) axon |
| $G, G_k^{axoplasm}, G_{m,k}$ | conductance (axoplasmic, myelin) of the ($k$-th) axon |
| $\theta_k, \theta_k(z)$ | ($z$-dependent) inclination of the $k$-th axon with respect to the central axis of the containing cylinder |
| $W_{kp}, W_{kp}(z)$ | a ($z$-dependent) matrix that contains the inter-axonal distances |
| $K_p, K_p(z)$ | a ($z$-dependent) parameter related to the $p$-th axon |
| $I, I_k^{injected}$ | the current injected, such as ionic (or stimulation) current, into the ($k$-th) axon |
| $I_k^{fields}$ | the field-generated current |
| $\vec{E_{1->2}}, \vec{E_{2->1}}$ | electric field generated by axon 1 (or 2) impacting on axon 2 (or 1) |
| $\vec{p_i}$ | the dipole moment of all nodes on axon $i$, superposed |
| $l$ | mean distance of all nodes of axon 2 from the present node on axon 1 |
| $\vec{l_i}$ | the vector containing all the distances relative to axon $i$ |
| $l_{ij}$ | the distance between the present node on axon $i$ and node $j$ on the other axon |
| $\vec{p_i^j}$ | the dipole moment of node $j$ on axon $i$ |
| $C$ | capacitance of the axonal membrane |
| $\eta$ | a constant appearing in the Hodgkin-Huxley equation |
| $\tilde{W}_{kp}(z)$ | ($k, p$)-th entry in the ($z$-dependent) modified $W$ matrix, as specified in this paper's text |
| $N$ | total number of axons in the axon tract |
| $\vec{\tilde{W}_k(z)}$ | ($z$-dependent) $k$-th row of the modified $W$ matrix |
| $\vec{V(z, t)}$ | ($z, t$)-dependent column vector containing the transmembrane voltages of all the $N$ axons |
| $u_k$ | linear combiner output of the $k$-th neuron |
| $w_{kj}$ | synaptic weights of neuron $k$ |
| $m$ | total number of synapses in a neuron |
| $x_j$ | input signals to the synapses of a neuron |
| $\varphi(\cdot)$ | activation function of the neuron |
| $b_k$ | bias of the $k$-th neuron |
| $y_k$ | output signal of the $k$-th neuron |
| $T_1, T_2$ | axon tract (1,2) |
| $SW_{T_j}^i(z)$ | subspace $i$ of the ($z$-dependent) $W$ matrix of axon tract $j$ |
| $F$ | a matrix, akin to the $W$ matrix, but for *collections* of tracts; the $W$ matrices ...<br>... of the component tracts are sub-matrices of this larger matrix |
| $o$ | a type of operation within a $W$ matrix |
| $s$ | a type of operation within an $F$ matrix |

This equation models nodes and internodes. In this equation, the capacitance enters on the left hand side. In order to model a paranodal segment, we modify the value of *c_m,k_*. This is discussed in detail in Section 3.1.

When Equation (1) was introduced, the matrix containing the interaxonal distances, *W_kp_*, was phenomenologically placed. Here we present a justification of the introduction of this matrix. The justification is based on electric field based coupling between the axons. In future work we will explore a purely current-based justification. We focus on the term 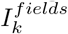 which stands for the current injected at a node of axon *k* based on the impinging electric field from other axons’ nodes. In particular, consider the electric field generated by axon 2 in its impact on axon 1, which enters into Equation (1) as a product with the conductivity in order to form the field-generated current injection at a node. Denote it by 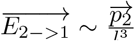 where 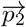 is the dipole moment of all nodes on axon 2, superposed (see Equation 4.13 in [12]), and *l* is the mean distance of all nodes of axon 2 from the present node on axon 1 (the node under consideration). Next suppose we denote by 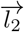 the vector containing all the distances: 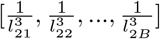 and by 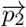 the vector containing all the dipole moments: 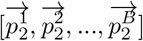. Then 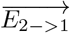 is proportional to the inner product between these two vectors. Similarly 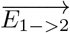 is proportional to the inner product between 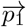 and 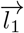. If we collect all these inner products in a given axon’s equation, they lead to a cubic-decay *W_kp_* matrix.

### 2.2 Demonstration of the Program

The basic program was outlined in a flowchart in [13]. In this section we present a numerical experiment that demonstrates the working of our program, extended as follows. In order to demonstrate the power and versatility of the program, we extended it to include Gaussian random noise at the ion channels (this was also done in [14], but for shorter spike trains) to introduce randomness into the simulations. We start with a tract of *N* = 3 axons and do not invoke electric fields or focal demyelination. We stimulate the first axon (topmost sub-figure in both Figures 1 and 2.). We record the voltage from the stimulated node on all the three axons as well as from the terminal nodes in all three axons. The stimulation magnitude is approximately 1.6e-5 Amps/centimeter. To achieve this, we inject 4 nano Amps in a node of Ranvier which has a length of 2.5e-4 centimeters. For biological comparison, the typical voltage gated sodium ion channel current is of the order of 2 pico Amperes, but several thousands of these are present at a node and superpose in some way to generate an action potential-like current profile. In [9], the authors inject a similar amount of current. In the present studies, the current is injected in short square pulses of duration 10 microseconds each, input at node 20 at a frequency of 1 per 2 milliseconds. Note that the duration of an action potential from our simulation program is about 1 millisecond, but if we include the refractory period, we cannot have action potentials less than about 2 millisecond apart. From Figure 1 the 2 millisecond choice is just about right, though a slightly larger temporal gap might have allowed equal magnitude of all the action potentials; the second and third action potentials for example appear slightly diminished in amplitude, which would be consistent with having an overlap between the refractory period of the preceding action potential and the injection of the present current pulse. The results are shown in Figures 1 and 2. The figures show that random ion channel noise can lead to fluctuations in the voltage recorded.

**Figure 1:**
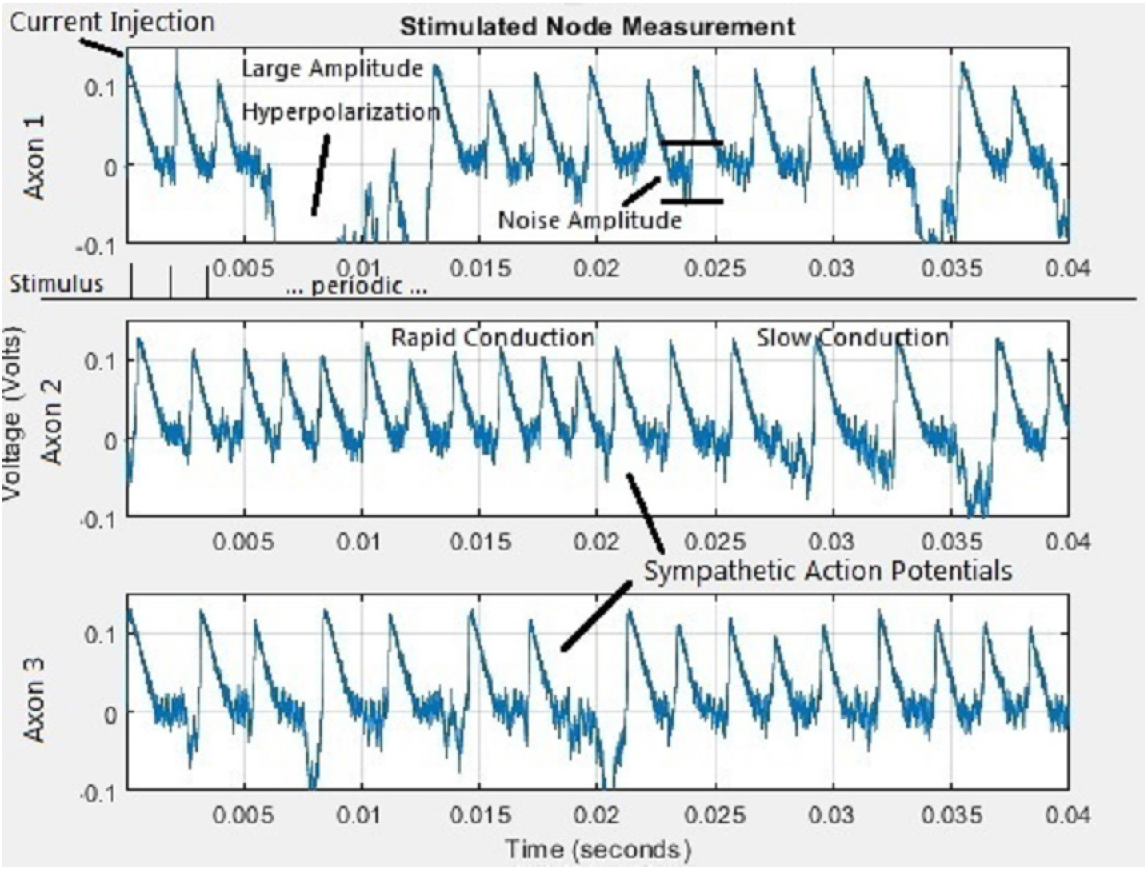
Voltage probe at the stimulated nodal position. Only axon 1 is stimulated at position 20, but all three are shown, with axon 1 the topmost sub figure and axon 3 the bottom subfigure. The *x*-axis is time in seconds and the *y*-axis is voltage in volts, in all the three subfigures. The severity of the added noise causes occassional large amplitude hyperpolarization.

**Figure 2:**
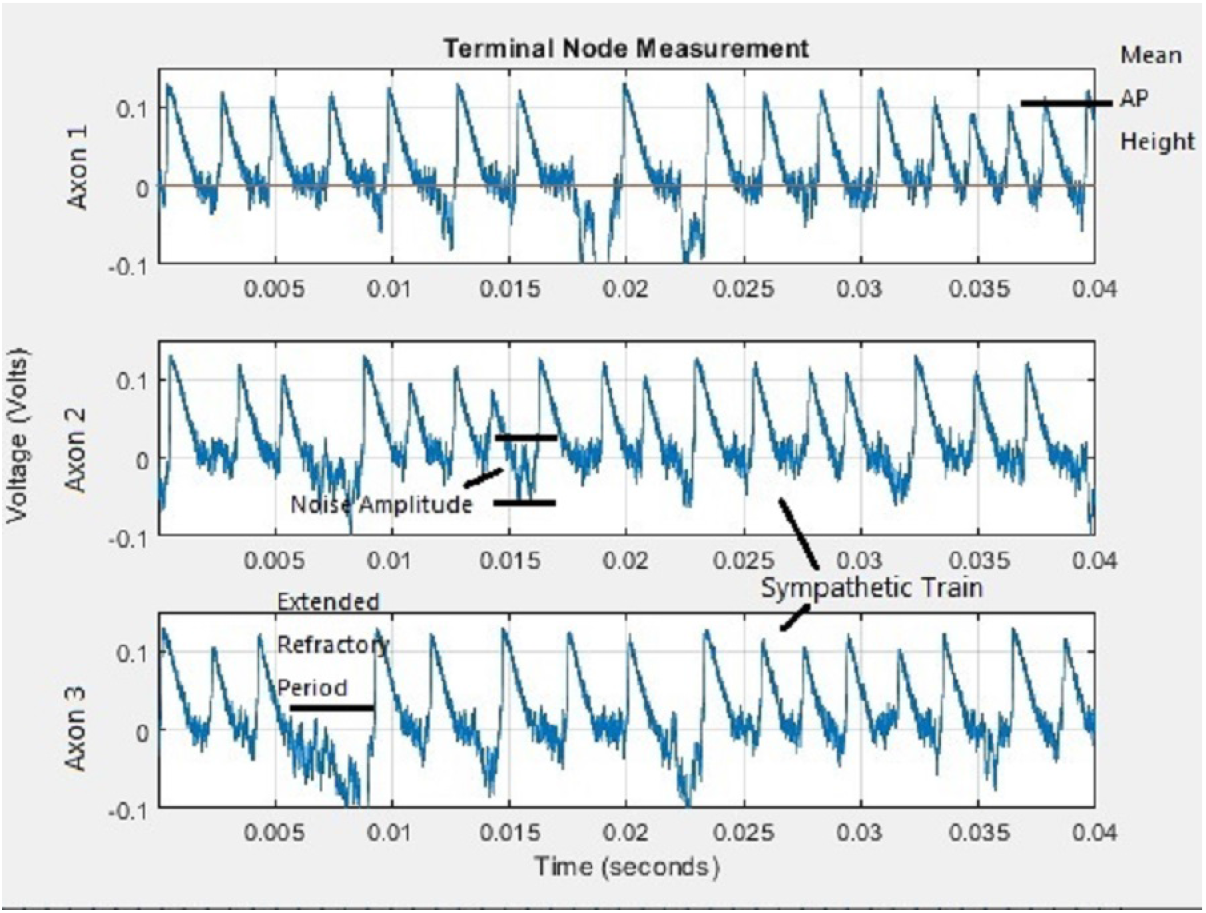
Voltage probe at the terminal position of all three axons. All three are shown, with axon 1 the topmost sub figure and axon 3 the bottom subfigure. The *x*-axis is time in seconds and the *y*-axis is voltage in volts, in all the three subfigures. Propagation delays are clearly visible in the first and second axons (top two sub-figures). The third axon does not show significant propagation delay; the explanation may lie in a combination of noise and current-coupling between the axons.

### 2.3 Theoretical and Geometric Analysis of Synchronization

In this section, we will present an analysis of synchronization from a geometric perspective. We will model the signal space geometrically after introducing an inner product on it. This will serve to illustrate a key concept of synchronization. Consider the following equation:

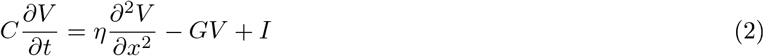

This is the (Hodgkin-Huxley) equation for a single axon. From [11], we note down the equation for a tortuous tract of N coupled axons:

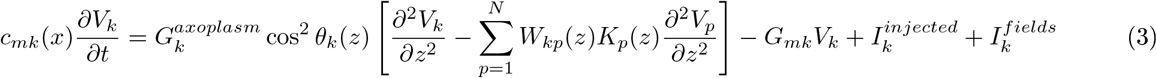

Introduce

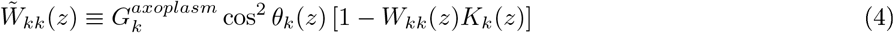

and

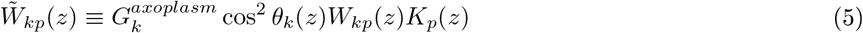

This 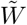 matrix has different structure on the diagonal as opposed to the off-diagonal. Inserting Equations (4) and (5) into Equation (3) above, we obtain,

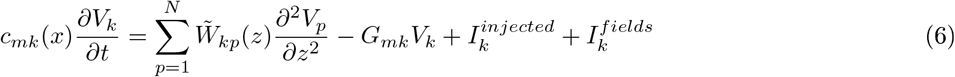

The summation sign in this equation can now be treated as an inner product. Explicitly we note,

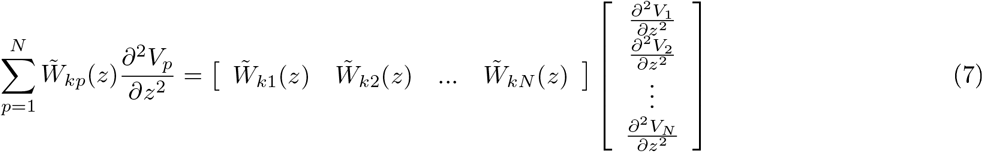

Henceforth we will denote the inner product in this formula as,

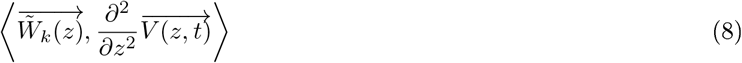

Inserting this back into Equation (6) we obtain,

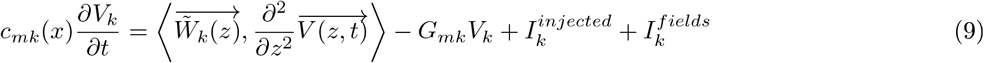

Note that 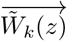 can be thought of as the geometrical “view” from the k-th axon and 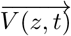 is nothing but the vector of transmembrane voltages of all the axons at the pair (*z, t*). In this form, Equation (9) tells us that the temporal evolution of the k-th axon’s voltage depends upon (in addition to the last three terms) the projection of the system’s voltages upon the view from that axon. Since the inner product is known to be related to the geometrical distance between the constituent vectors, this gives us the following picture of what is happening with time (see also Figure 3).

**Figure 3:**
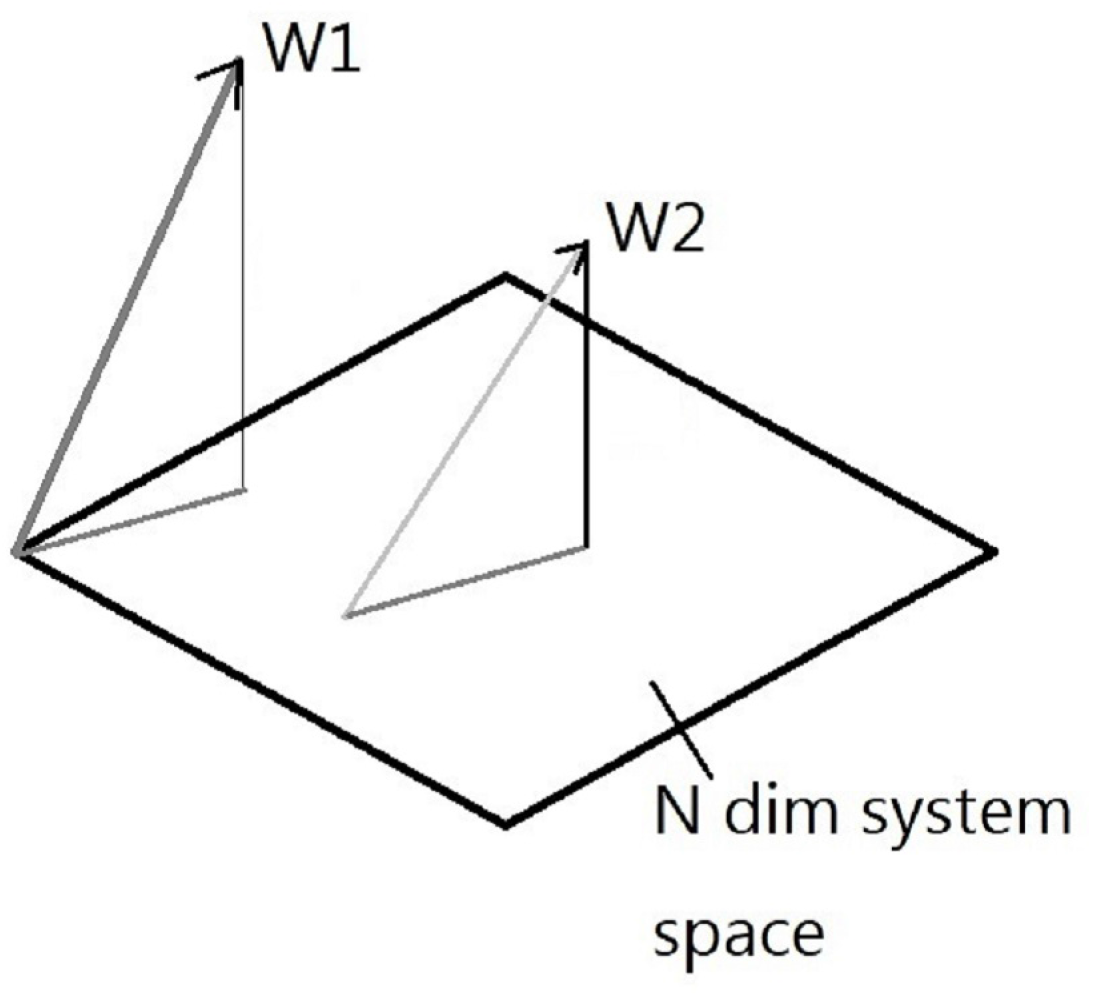
Geometrical view of the evolution equation (9). Following formula (8), the *W*-vectors (*W*_1_,*W*_2_ and others) are projected (via inner products), onto the *N*-dimensional system space comprising the span of the voltage vectors whose entries are *V*_1_ through *V_N_*. These *N* voltages, in this format, comprise a sub-set of ℛ_N_ but, by using the so-called *push-forward construction*, can also be viewed as a subspace of ℛ_N_.

We start off the system with some set of voltages on all the axons and some fixed geometry. As time progresses, each voltage migrates according to the law in Equation (9). The migration rate of a voltage is higher if its separation (by which we mean the separation of its geometrical vector) from the system voltage vector is larger (modulo a linear term). Visualize the system voltage vector as a cloud (see also Figure 4). If any point in the cloud is far from the cloud (in a mean sense), then the point will evolve at a much more rapid rate, than points closer to the cloud. If this evolution is towards the cloud, it will result in synchronization. If the evolution is away from the cloud then it will result in dispersal of the voltages. Since we know that synchronization *does* take place in such systems, it must be the former case. Thus we have a geometrical view of synchronization in axon tracts.

**Figure 4:**
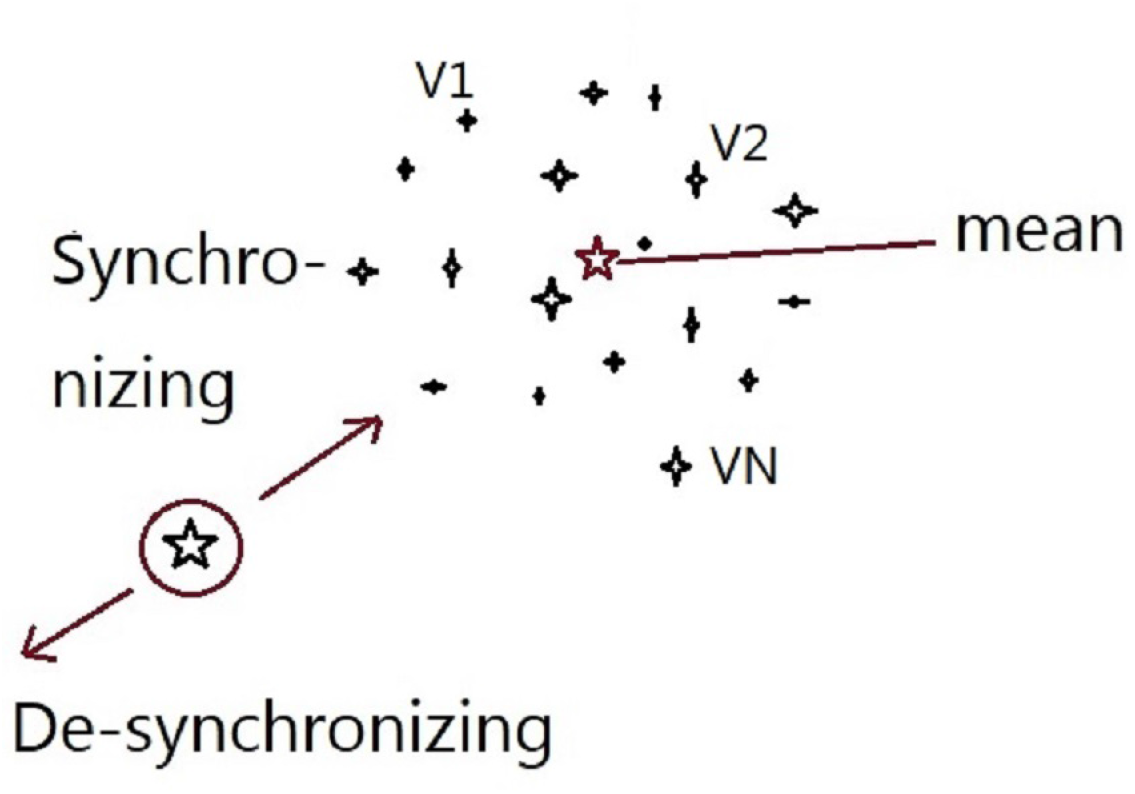
Cloud of points representing each of the system’s *N* transmembrane voltages. A distant point (shown circled in red) can evolve over time to approach the cloud or recede from it. If it does the former, the point is said to be synchronizing with the rest of the system and in the latter case it is going away from the mean.

### 2.4 The Tract as a Learning Machine?

In this section, we present the ephaptic equation of [13] as an artificial neural network (ANN) with feedback with the aim of showing that an axon tract can learn in the sense of machine learning. As per [15], a neuron, labeled *k*, may be described by writing the pair of equations:

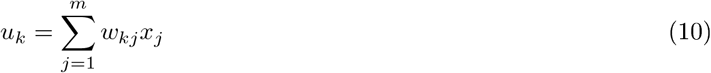

and

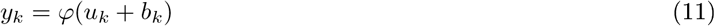

where *x*_1_, *x*_2_, *x*_3_, . . ., *x_m_* are the input signals; *w_k_*_1_, *w_k_*_2_, *w_k_*_3_, . . ., *w_km_* are the respective synaptic weights of neuron *k*; *u_k_* is the linear combiner output due to the input signals; *b_k_* is the bias; *φ*() is the activation function; and *y_k_* is the output signal of the neuron. [15] further says that the bias *b_k_* is an external parameter of neuron *k*; and we can reformulate the model of neuron *k* by doing two things:

1. adding a new input signal fixed at +1, and
2. adding a new synaptic weight equal to the bias *b_k_*, resulting in a total of *m* + 1 synaptic weights instead of the *m* original ones before.

Recall the central equation from [13] as being given by (Equation (1)):

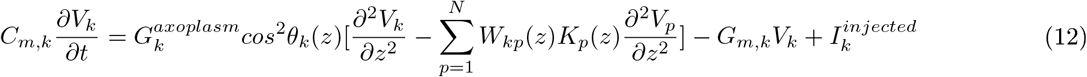

where *k* = 1, . . ., *N*. We focus on the nodal equation and expand the parentheses on the right hand side (RHS) to obtain:

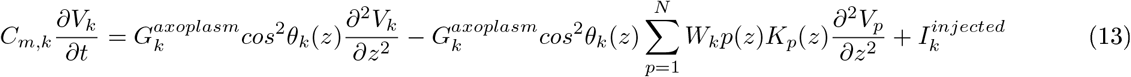

where *k* = 1, . . ., *N*. Next we specialize to the *k* = 2 case and obtain:

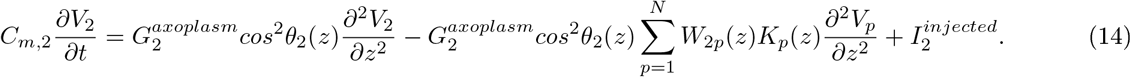

We group all the explicit *V*_2_ terms on the right hand side together (ignoring the injected current for the moment) to obtain:

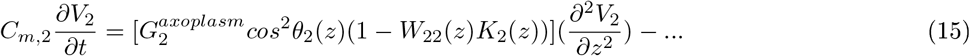

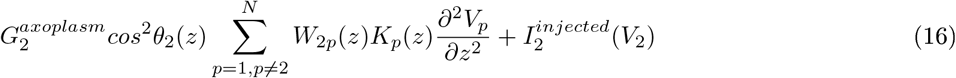

Since the derivative of a sum of functions distributes over the functions, we take the derivative out of the summation on the right hand side to obtain:

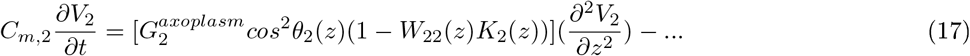

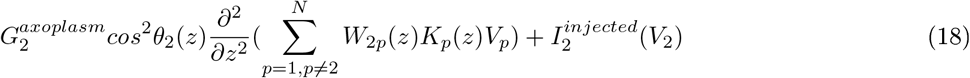

Next, we look at the injected current term which we previously ignored. Using the Fundamental Theorem of Calculus [16], we can rewrite the last term on the right hand side as

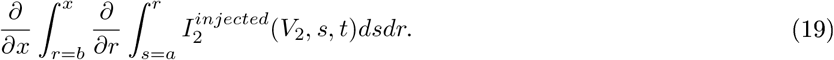

The conditions when we can write functions as second partials of double integrals [see Theorem 9.42 in [17]] need to be further examined in future work. Nevertheless, if we continue along the lines presented in the last three equations above, we will end up with a formulation like Equation (11), with weights and bias terms on the right hand side and the voltage of the *k*-th axon on the left hand side - having defined a suitable activation function. Interpreting the formulation as suggesting a perceptron with feedback, we might be able to infer that *an axon tract can learn*.

## 3 Results

As mentioned in Section 2, in [13] we introduced a MATLAB program (henceforth referred to as M2020) for simulating three-dimensional axon tracts. Those tracts did not have demyelination - that is, there was a uniform myelin coverage on all the internodal regions. However, in diseases such as multiple sclerosis, the myelin coverage is no longer uniform. In this section we will approach the problem of simulating axon tracts when the individual axons can have spot-demyelination, also known as ‘focal’ demyelination, along their length.

### 3.1 Results From Modification to MATLAB program M2020

These experiments indicate that sodium channels form along the internodal segments of demyelinated axons [18].

The above statement from [18] motivates us further to introduce focal demyelination in internodal segments. Focal demyelination is simulated by creating a third type of segment, known as a paranode, apart from the nodal and internodal types. The only difference between a node and a paranode is in the capacitance per unit length [19]. If the capacitance is high, the rate of flow of charge, or current into the capacitor will be high. As a result, by Kirchhoff’s nodal law, the current in the conductance will be lower. By Ohm’s law, this would imply that the potential drop across the conductance, will be lower. As a result the action potential will reach a lower height than when the capacitance is low.

This phenomenon is observed at the paranodes, where capacitance per unit length is higher than that of the nodes. As a result, the action potential reaches a lower height at the paranodes than at the nodes. For a fixed current, the rate of rise of the action potential will be lower for a higher capacitance. Thus the rate of rise of the action potential at the paranodes is slower. The rate of diffusion across the succeeding internode to the next node is also slower and so the action potential train slows down after passing through the paranodes. This is also observed and verified as shown in Figure 5 below.

**Figure 5:**
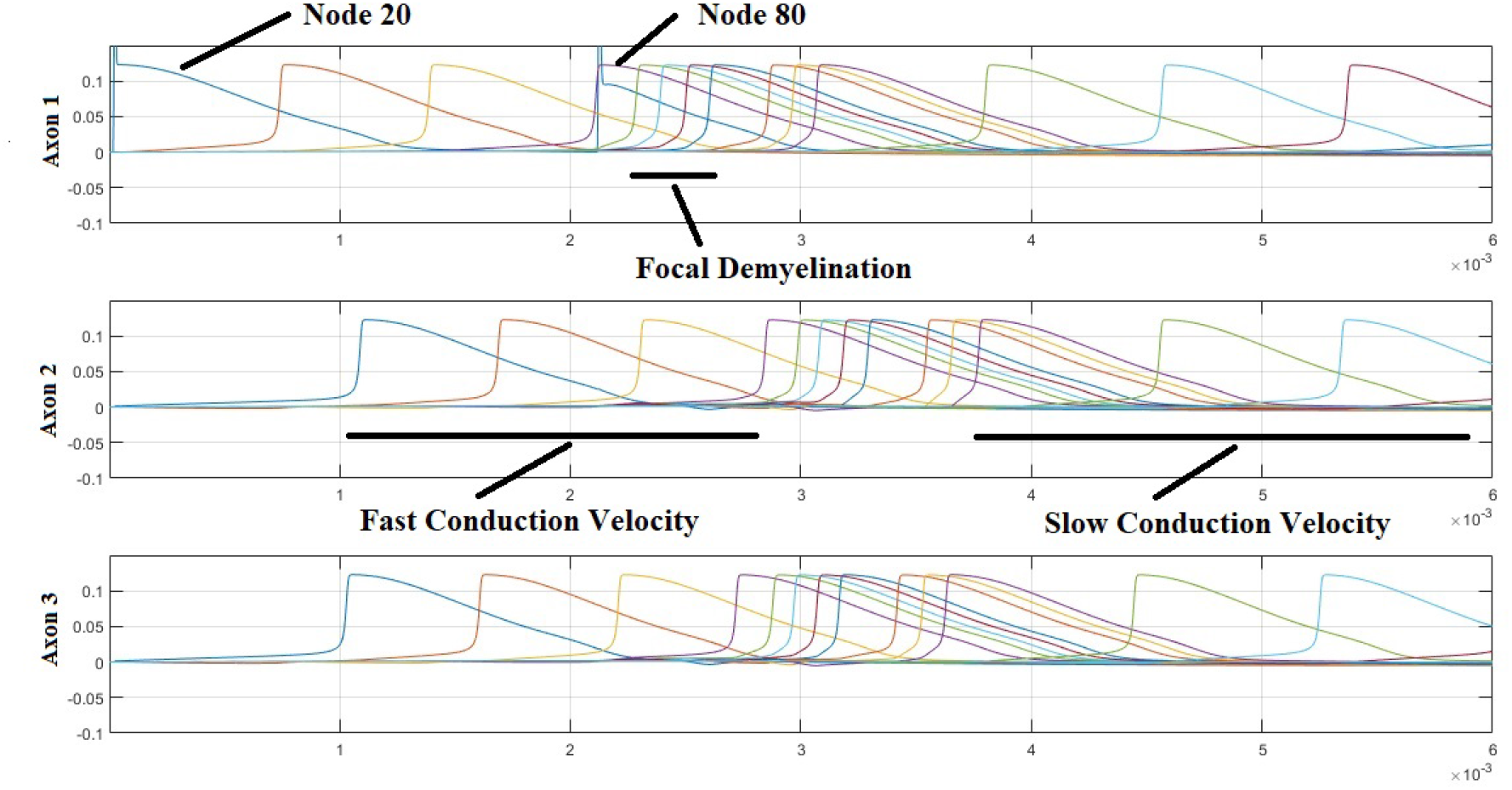
Slower and lower: Transformation of the action potential train on encountering paranodal regions with higher capacitance per unit length. The *x*-axis is time in seconds. The *y*-axis is voltage in volts.

At this stage, we would like to make a quick note that the studies in ([20], Figure 1A) indicate that axon tracts may demyelinate in various geometries within the tract cross-section, and there is indication that these demyelinated regions will contain axons with positions of demyelination that are all on the same ‘level’ - the level of the cross-section taken. Thus axonal misalignment might not play a particularly significant role in the placement of the paranodes, within the biological-experimental context. Therefore, in our simulations in this paper, we placed paranodes ‘facing’ each other on various axons in a tract, instead of ‘staggering’ them.

### 3.2 Variation of Conduction Delays With Length of Internodal Regions

We vary the length of the internodes and measure the resultant change in conduction velocity. This is done in order to verify the remarks made in [21] indicating absence of continuous conduction in fibers with long internodes. In Figure 5 it is clear that the initial conduction velocity is 2.7273 meters per second while after passing through the three paranodes, the conduction velocity is about 2.3529 meters per second (a 13.73 percent change). In the next figure, Figure 6 below, we increase the length of the internodal regions by a factor of two. We observe that conduction stops completely after a single post-paranodal firing.

**Figure 6:**
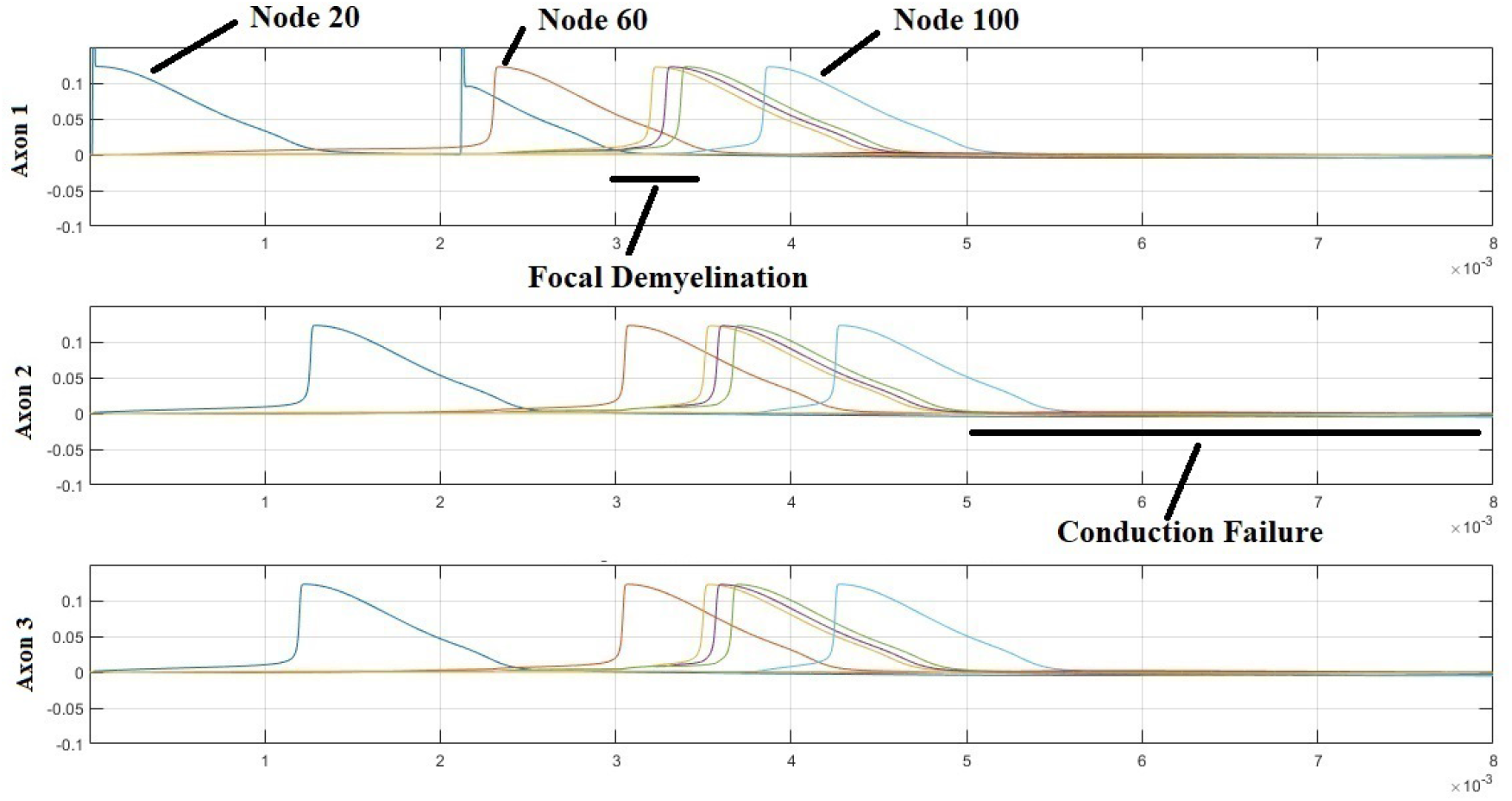
An action potential train encounters a region of focal demyelination and exhibits post-paranodal conduction failure. The *x*-axis is time in seconds and the *y*-axis is voltage in volts.

That Figure 6 does not simply represent a much larger conduction delay was verified by zooming in on the next node after the last node with a bonafide action potential. The following figure, Figure 7, shows the result of zooming in. It is clear that the peak voltage attained is less than 2 mV, which is fairly below threshold.

**Figure 7:**
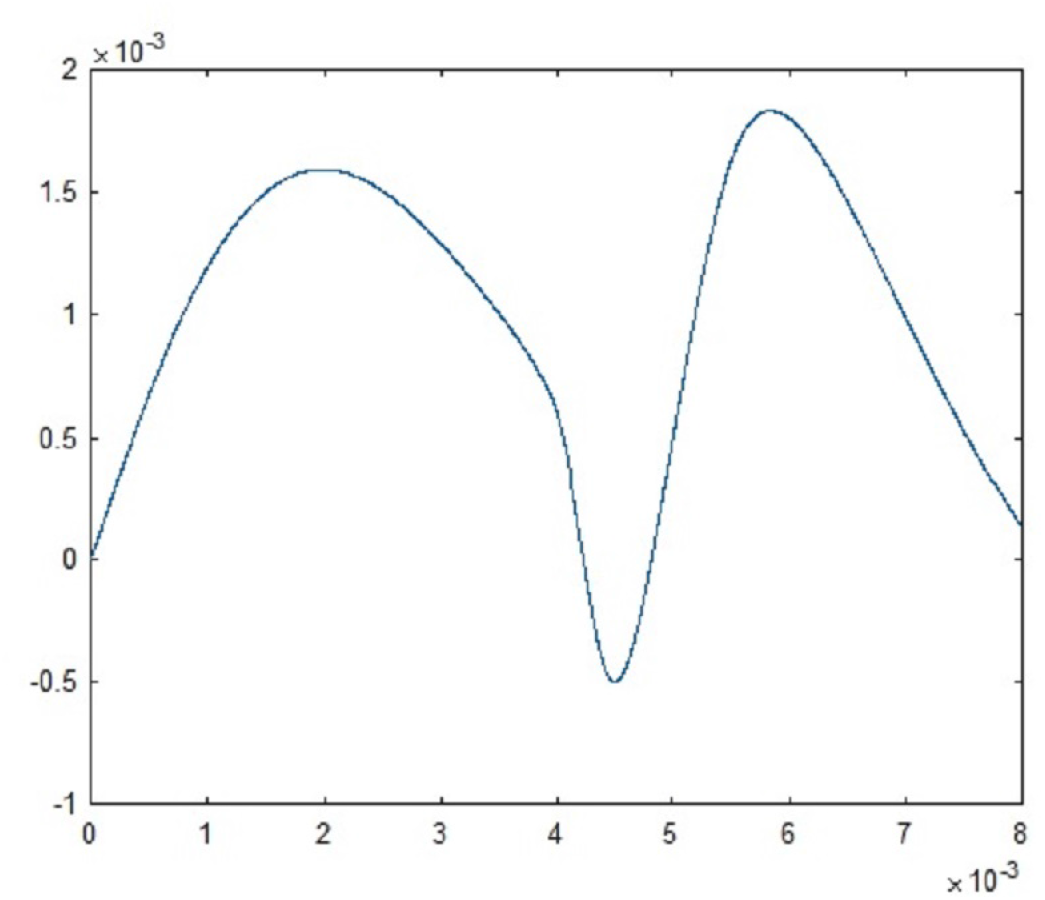
Subthreshold excitation on the next node (node 140) after the last excited node (node 100), for the case simulated in Figure 6. The *x*-axis is time in seconds and the *y*-axis is voltage in volts.

In Figure 7, the paranodal capacitance has been set at 5.14*e*−9 F/cm (within 2*e*−9 F/cm of the nodal capacitance). Next, we wish to ascertain if the conduction failure is aided by this high value of the paranodal capacitance. Towards this end, we obtain the next figure, Figure 8, which shows that even with a lowered value of paranodal capacitance, 4.14*e* − 9 F/cm, there is conduction failure (see also [13] for the normal capacitance value). Additionally, we note that the final action potentials on the three axons have a modified inter-action potential temporal separation as well. Thus the data that is output after the paranodes is also modified (say, as in the case of time coding) with reference to that before reaching the paranodes.

**Figure 8:**
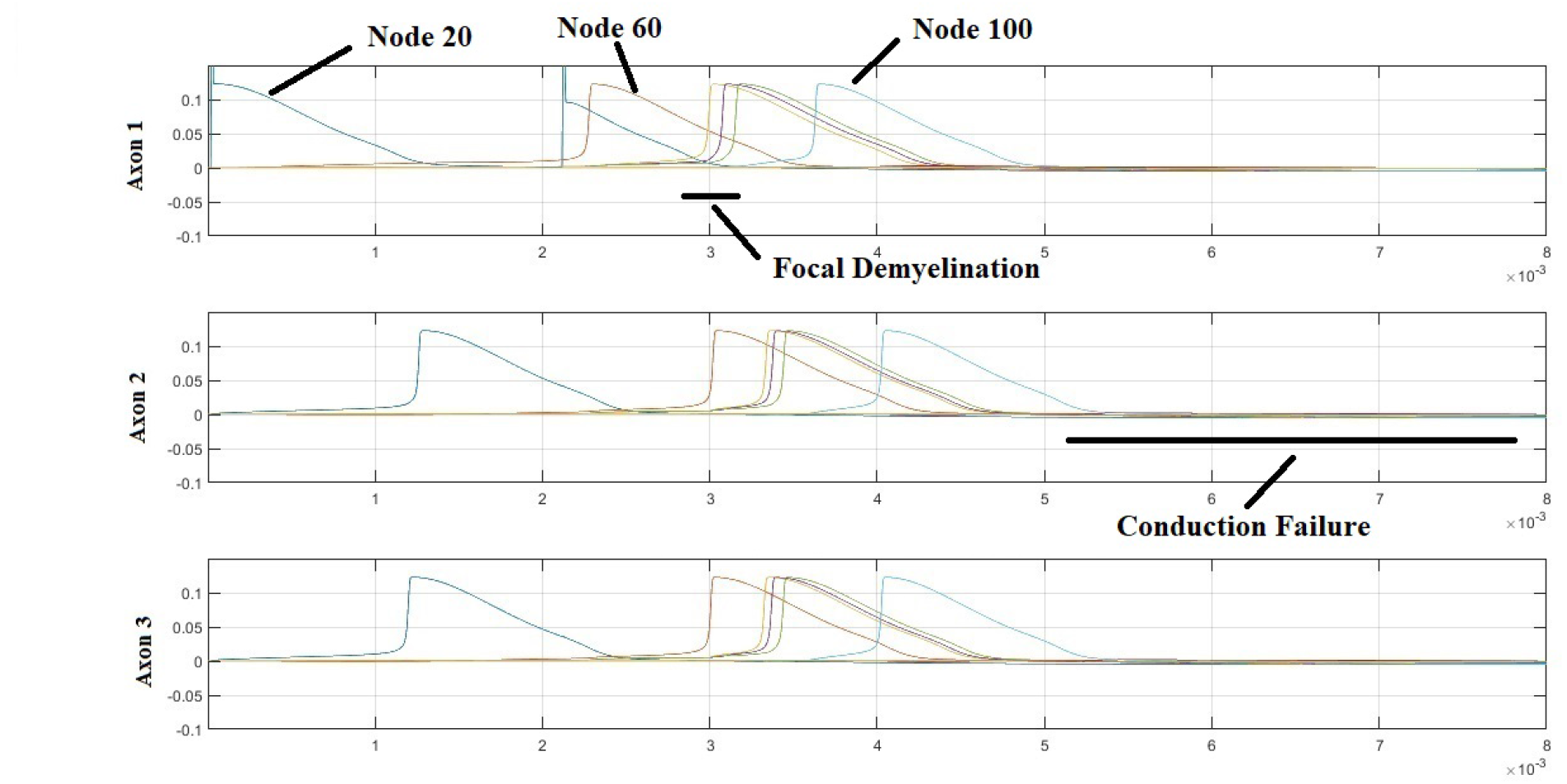
Conduction block even with a lower paranodal capacitance (4.14*e* − 9 F/cm) as compared to the previous case, Figure 6. Modified temporal separation ratios between the three axons’ action potentials post paranodal transit, indicating modification of time coded data. The *x*-axis is time in seconds and the *y*-axis is voltage in volts.

Finally, a quick note on trying to replicate the slowing down of action potentials of Figure 5 without careful thought; all we get initially is a conduction block. If, however, we recall that the internodal spacing used for generating Figure 5 is 20 as opposed to 40+, then, when we reduce the spacing used in the replication down to 20, we right away obtain the same slowing down as observed in Figure 5. This phenomenon is related to the interaction between conduction velocity and internodal spacing [22].

### 3.3 Electric Fields and Focal Demyelination

In this section, we carry out the investigations of the previous subsections, but with electric field ‘gain’ turned on. This enables the fields generated at the nodes to influence the ions at distant nodes. However, just like in the program presented in [11] (M2021), electric fields are not computed using actual node-to-node distances, but only approximate distances. We also make the simplifying assumption that paranodal fields cannot influence other nodes and paranodes.

When we carry out the simulation just described, we obtain the results shown in Figure 9. These results assume equal field ‘gains’ on both the nodes and paranodes. The field gain is a multiplicative coefficient which can be set to zero to turn off the contribution of the corresponding fields, and to a positive real number to turn it on. By setting the gains to be equal, we treated the nodes and the paranodes on an equal footing, in field-contribution terms.

**Figure 9:**
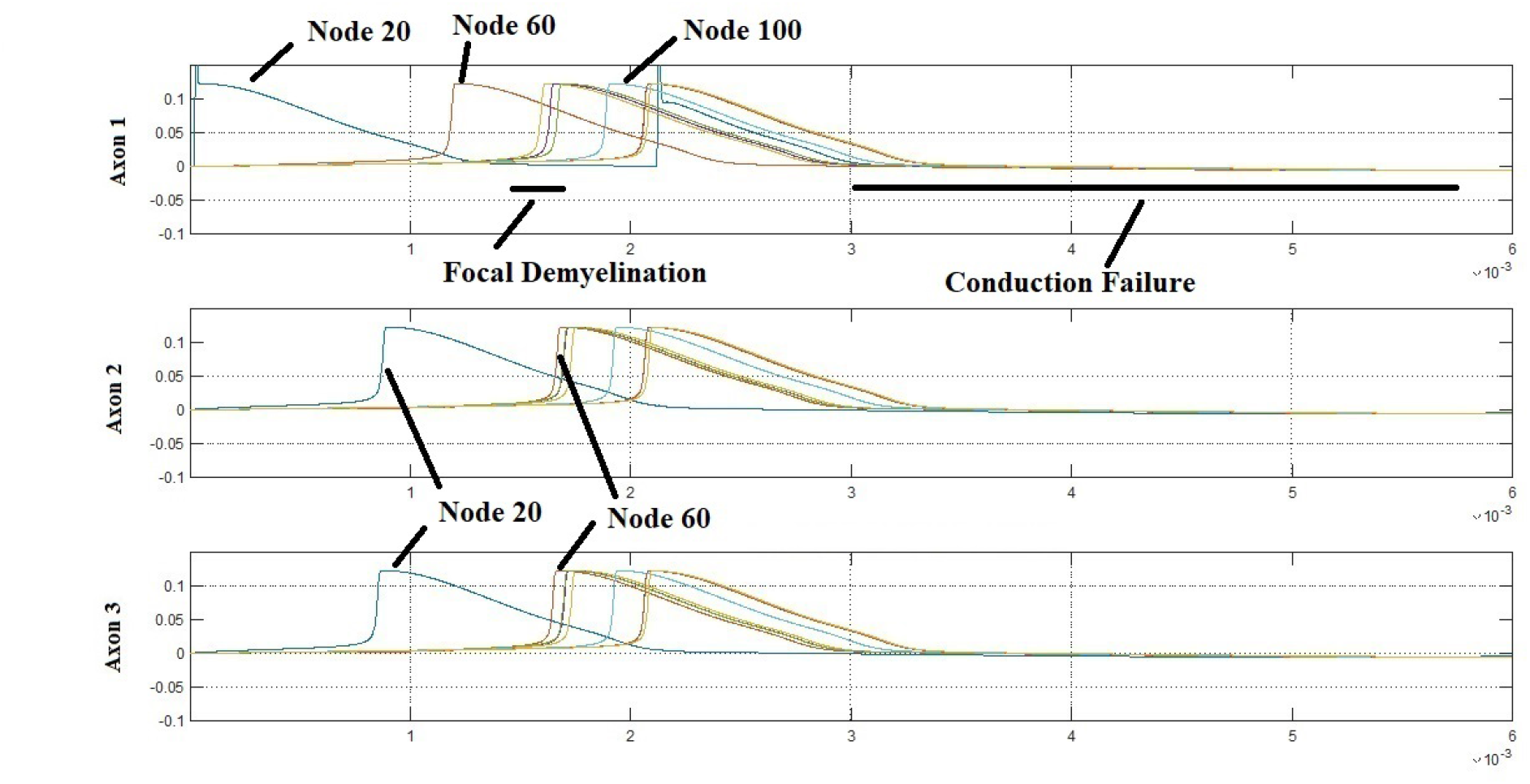
Focal demyelination in the presence of electric fields, with equal field gains of 5*e* − 13 for the nodes and paranodes is here simulated. There were three paranodes 84, 86 and 88 and a nodal spacing of 40, yielding nodes at positions 20, 60, 100, 140, 180, 220, 260. Conduction block is clearly observed. Also note that there are three more action potentials after the paranodes are crossed. Further note that the conduction block happens synchronously on the three axons. The *x*-axis is time in seconds and the *y*-axis is voltage in volts.

### 3.4 Catch and Re-inject Simulations of Tortuous Tracts

Tortuous tracts were introduced in [11] along with their governing equation. Figure 10 below shows a schematic of a tortuous tract with two segments. The first segment has axonal inclinations of *a* degrees and the second one of *b* degrees, imparting a tortuous nature to the overall tract.

**Figure 10:**
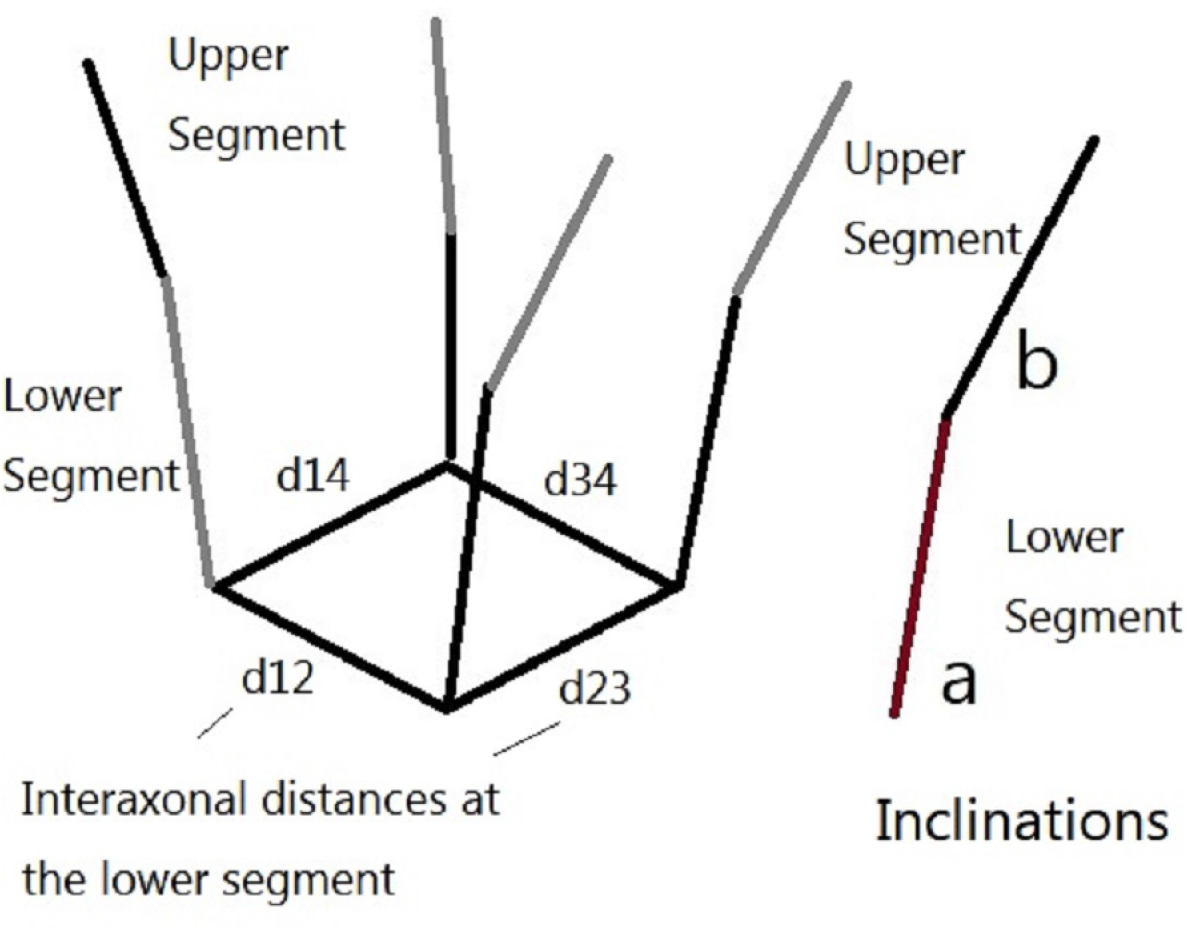
Schematic of tortuosity. *a* and *b* are the inclinations of the axons in segments 1 (lower) and 2 (upper) respectively.

We use a catch and re-inject method in order to simulate the impulse propagation in a tortuous tract whose three successive segments have axons inclined at 50, 55 and 60 degrees respectively. In this method, we make short segments of the first inclination. Upon injecting this segment, the output action potentials are ‘caught.’ These are then used to determine the stimulation on the next segment. This is known as ‘re-injection.’ These segments are stacked together back-to-back in order to form the tortuous tract. It is clear from Figures 11 and 12 that the action potential train has slowed down as a consequence of the tortuosity. To validate this further the next two figures, Figure 13 and Figure 14, show a non-tortuous tract of the same number of segments and inclinations 50, 50 and 50 degrees each. These observations have consequences for our understanding of why tortuosity is prevalent in the higher brain regions, but not as pronounced in the peripheral nervous system.

**Figure 11:**
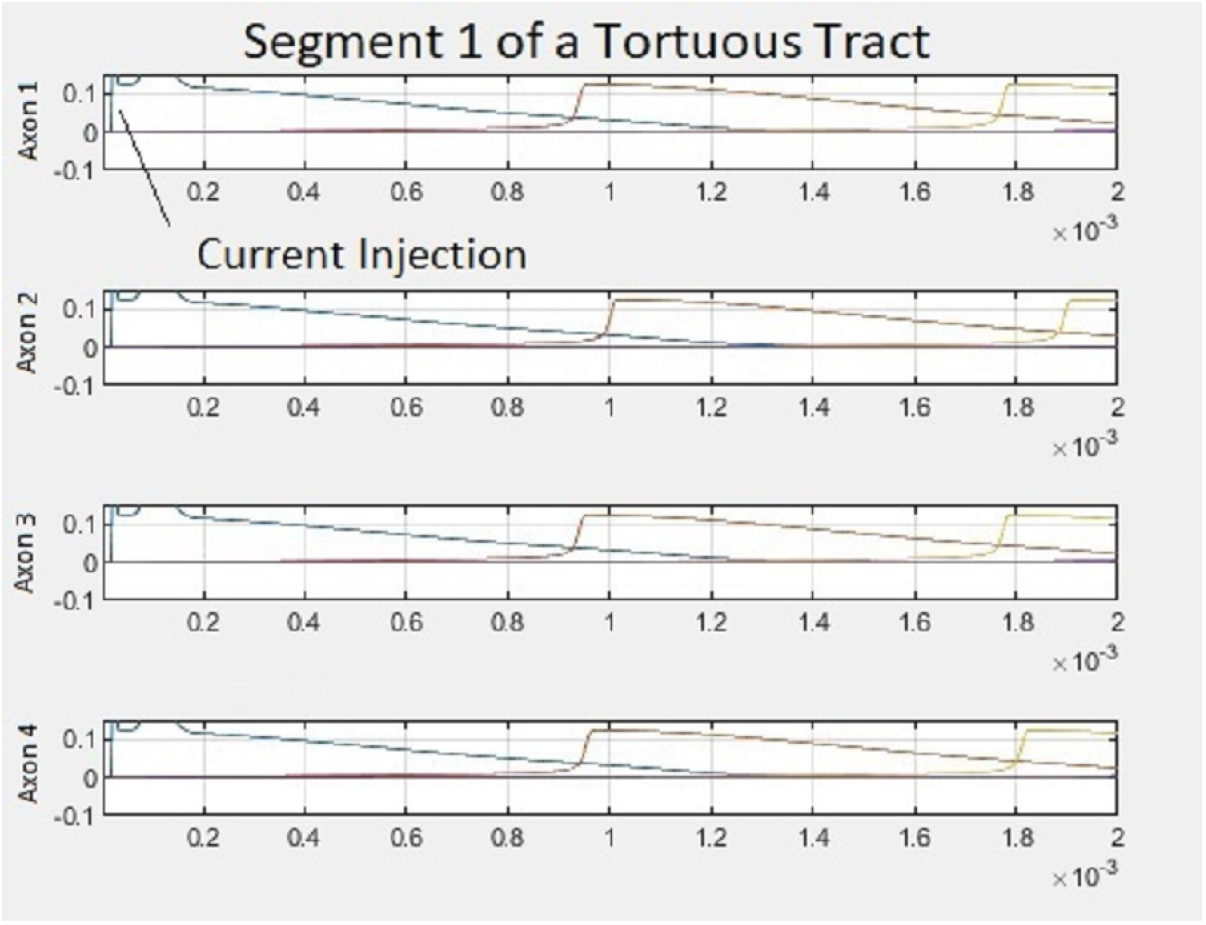
The initial segment of a tortuous tract. Note that the action potentials are about 3 per subplot. This tract segment has axons inclined at 50 degrees to the tract axis. The *x*-axis is time in seconds and the *y*-axis is voltage in volts, in every sub-plot. The interaxonal distances are not symmetric and are a likely cause for the difference in propagation speeds on the four axons.

**Figure 12:**
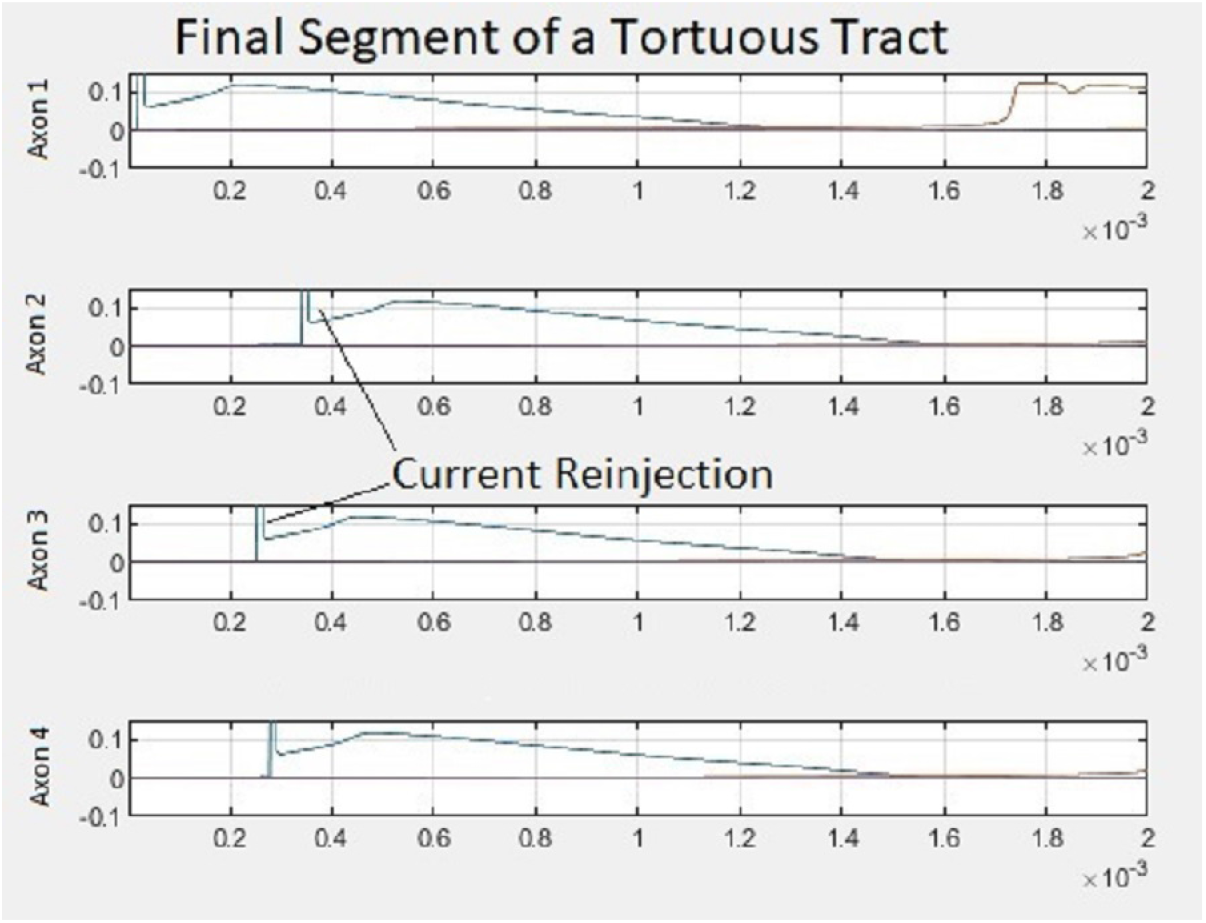
Final segment of the tortuous tract shown in Figure 11. Note that roughly 2 (or fewer) action potentials are visible per sub-plot, indicating a relative slowing down of the impulse train. This tract segment has axons inclined at 60 degrees to the tract axis. The *x*-axis is time in seconds and the *y*-axis is voltage in volts, in every sub-plot.

**Figure 13:**
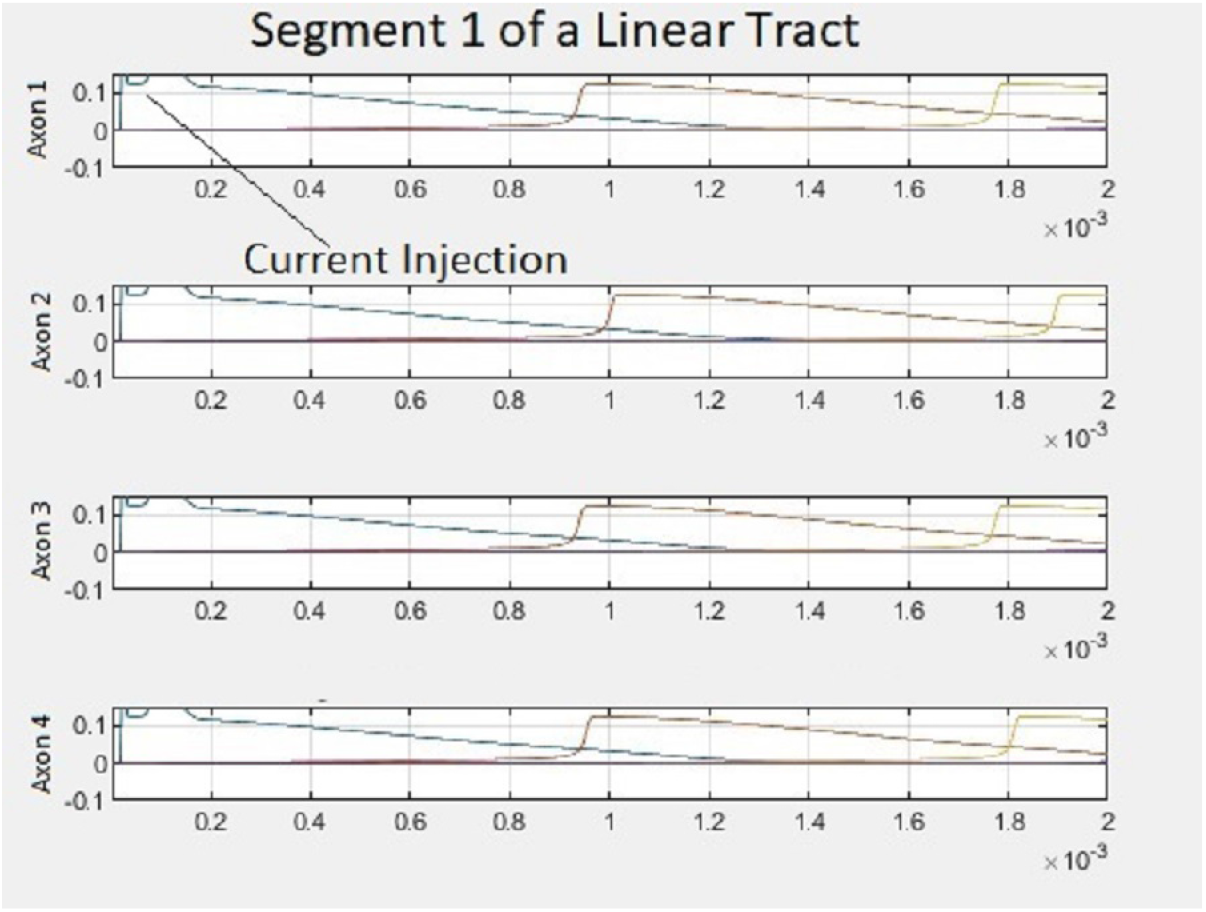
The initial segment of a tract with equal inclinations in each segment. In this segment, the inclination is 50 degrees to the tract axis, just as in Figure 11. The *x*-axis is time in seconds and the *y*-axis is voltage in volts, in every sub-plot.

**Figure 14:**
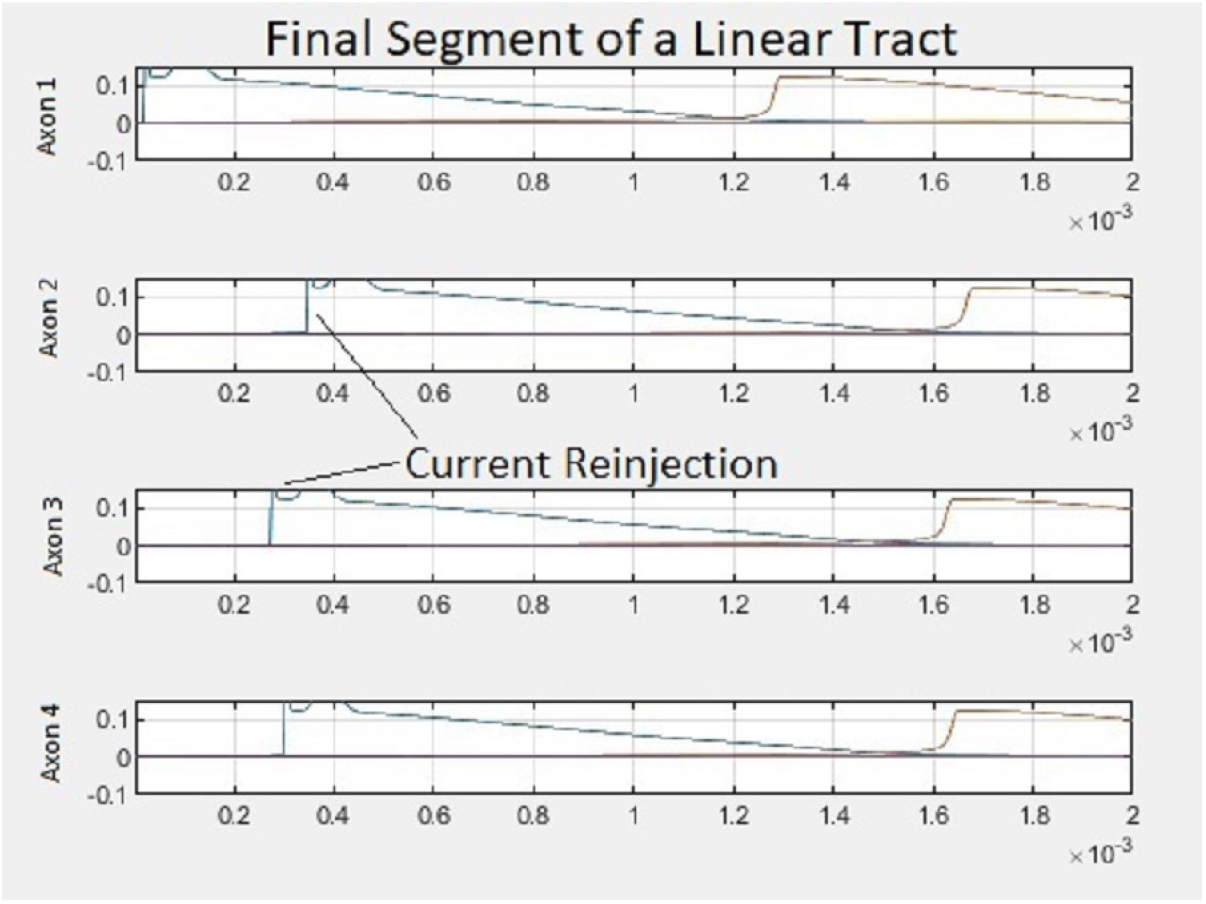
Final segment of a linear tract with axonal inclinations of 50 degrees in each segment. It is clear that conduction velocity in the third segment is higher for the linear tract, as compared with the tortuous tract of Figure 12. The *x*-axis is time in seconds and the *y*-axis is voltage in volts, in every sub-plot.

## 4 Discussion and Conclusion

Studying the conduction velocity in axons is clearly important from both mathematical and clinical viewpoints. As discussed in our forthcoming (2022) paper, the most relevant way of deducing the conduction velocity from computer simulations is to find its value by determining the inter-spike interval, and using it in conjunction with the inter-nodal spacing. We used this methodology in our study of focal demyelination. First, we looked at the case where the axons in an axon tract are current-coupled. We found that focal demyelination could induce either a conduction block (failure) or lead to a changed post-demyelinated zone conduction velocity. Next we looked at demyelination in the presence of electric field mediated-coupling (in addition to current mediated-coupling). We found conduction block to occur again, but nearly synchronously on all the axons. This is consistent with the previously observed impact of electric fields on synchronization - they aid in rapid synchronization. We next discuss the functional purpose of rapid synchronization, placing our study in a broader neuroscientific context.

Why would we need rapid synchronization? This issue must be addressed if we are to take seriously the possibility that electric fields play an important role in axon-axon coupling. From the perspective of information processing, provided we are sure that synchronization aids information processing (see our 2014 poster), it is certainly beneficial (in some respect) to have the information processed rapidly. For instance, we can imagine signals that need to be at their destination ‘on time.’ In a fight-or-flight scenario, such a consideration might be relevant. However, there may also be reason for slow processing to take place - in fact, multiple timescales of information processing could be occurring simultaneously, some maintaining the slow diurnal clock for instance, and others processing faster events. Thus synchronization of all rates might be needed by the brain. A wide range of possible rates would afford ‘comfortable’ information processing across the spectrum of possible organismal needs. Thus some sort of mechanism could be searched for, or posited, which would allow a ‘weighted use’ of electric fields and electric currents - rapid and slow, as per the demands of the organism for a specific information transmission speed. We have not explored this question deeply in this paper, but what we can see is the different rates of synchronization in the presence of conduction block when fields were also included into the picture. This might have implications for the recovery and relapse periods of multiple sclerosis.

This study has enhanced our overall understanding of demyelination in the presence of current and field based axon-axon interaction, and in future work we intend to travel down two additional directions. The first will look at randomly placed zones of demyelination in a tract. The second will introduce channel capacity measurements as a quantifier of the demyelination, similar to the work done on measuring channel capacity under various geometries for healthy axon tracts in our 2021 paper. Furthermore, our justification of the use of the *W* matrix in our 2019 formulation is potentially quite significant when considering field-based interaction. Currents and fields are closely related and so we expect that in future work we will be able to provide a justification for the *W* matrix in the current-based interaction setting as well. With this, the mathematical model being developed since the publication of our 2019 formulation, is supported further. Our geometric picture of synchronization illuminates the mechanism of the synchronization process and is reminiscent of machine learning, especially unsupervised learning. Many concepts from machine learning may thus be relatable to the axon tract under coupling conditions and, in fact, we take the first rudimentary steps in showing that a tract may be viewed as a learning machine. These innovations provide new insights into axon tracts - the complexity of their function is aided by their geometric structure. Finally, in the appendix we make a brief note on how this geometry can also be viewed information theoretically - potentially providing a new paradigm in the study of diseased conditions.

## Appendix

## Appendix: A Note on an Algebra for W-matrices

In this appendix we discuss how two interacting nerve tracts may be represented by their *W* matrices [13] each, and how their interaction can be captured succinctly in an algebra for those *W* matrices. Once the tract has been represented as a learning machine [15] (see also Section 2.4), we have a unique matrix corresponding to the tract such that the tract ‘output’ (which is really fed back to the input) is this matrix times the input, if we ignore the bias.

Clearly, the *W*-matrix (along with the axonal inclinations) *W*_*T*_1__(*z*) influences how the axons within a tract *T*_1_ would interact via currents and/or fields. Might there be non-interacting subspaces in *W*_*T*_1__(*z*)? Note that here interaction refers to the ephaptic interaction which is current-mediated. Motivated by quantum mechanical decoherence-free subspaces [23], yet distinct from them, these non-interacting subspaces could be places in the tract where information would flow down the tract unimpeded by interaction. Hence if these subspaces 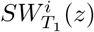 are found to be present even during advanced stage multiple sclerosis (MS), it would offer some hope to the patient’s clinician. If the tract is tortuous, the *W*-matrix is changing along the tract as indicated by the *z*-dependence. Thus these subspaces may pop in and out of existence with distance *z* along the tract. If we can find a connecting thread, with minimal interaction, linking the subspaces, then information could flow down this thread even in advanced MS. Any implantable medical neuro-device that we could design, should take advantage of this thread’s relative state of ‘protection’ from the disease. We may denote this thread as a disease-free information pipe (DIP). If two axon tracts or axon bundles *W*_*T*_1__(*z*) and *W*_*T*_2__(*z*) are juxtaposed, there may be no current-mediated interaction, but there could be field-mediated cross-talk that would couple the *W*-matrices. These *W*-matrices would then be thought of as sub-matrices of a larger *F*-matrix, within which we could again do an analysis similar to the one discussed earlier in the paragraph, but taking note of the difference - that the coupling is only field-mediated.

There may therefore be posited two types of operations, one type when considering sub-matrices within a given *W* matrix, denote this as *o*, and a second type when considering interactions between *W* matrices, or sub-matrices within a given *F*-matrix, denote this as *s*. Note that the operations are here said to be between sub-matrices or *W*-matrices. However, what we indicate by this is that they are between the transmembrane voltages of the axons in the tracts represented by these sub-matrices or *W*-matrices. Now, since each axon in *W*_*T*_1__(*z*) would interact with every other axon in *W*_*T*_2__(*z*), *F* might have a tensor product structure, i.e. *s* would be the tensor product operation. So finding DIPs in *F* would be more complicated than the procedure described in this appendix for the *W*-matrix.

